# Defective peripheral B cell tolerance leads to dysregulated B cell responses in Fibromyalgia Syndrome

**DOI:** 10.64898/2025.12.16.694591

**Authors:** Alexander Long, Antonio Choi Chiu, Orthi Onupom, Richard Berwick, Dimitra Psyllou, Jane Pernes, Katy Plant, Harvey Neiland, Andy Cross, Felicia Tucci, Andreas Goebel, Rachael Bashford-Rogers

## Abstract

Fibromyalgia syndrome (FMS) is a chronic pain disorder characterised by widespread musculoskeletal pain, fatigue, and cognitive dysfunction, with no definitive biomarkers or mechanism-based treatments. Emerging evidence suggests that immune dysregulation may contribute to the FMS pathogenesis, particularly involving B cells, which have been implicated in autoantibody production and neuronal sensitisation. However, whether peripheral B cell tolerance, a critical safeguard against autoimmunity, is compromised in FMS remains unknown. Here, we combined high-resolution B cell receptor (BCR) repertoire sequencing, deep immunophenotyping, and functional assays in a well-characterised FMS cohort to uncover profound defects in peripheral B cell tolerance. We reveal significant defects in peripheral B cell tolerance in FMS, including: (1) impaired naïve B cell anergy, marked by elevated CD21, CD22, and CD24 expression; (2) exaggerated proliferative responses and rapid CD24 downregulation upon stimulation; and (3) altered BCR selection patterns, with increased IGHV6-1/IGHJ6 usage, skewed class switching toward IGHA1, and enhanced heavy chain clonal expansion. These features closely resemble immune pathology profiles observed in classical autoimmune diseases. These findings redefine FMS as a disorder of immune dysregulation, with defective B cell tolerance contributing to disease mechanisms. The convergence of interferon-driven B cell activation, heavy chain clonal expansion, and autoantibody production suggests shared pathways with classical autoimmune diseases. Our study provides a foundation for mechanism-based diagnostics and targeted immunomodulatory therapies, offering new avenues for intervention in this debilitating condition.

## Introduction

Fibromyalgia syndrome (FMS) is a common chronic primary pain disorder characterised by widespread musculoskeletal pain, fatigue, cognitive dysfunction, and sleep disturbances ^1^. These symptoms significantly impair the quality of life, making FMS a debilitating condition for those affected. The prevalence of FMS varies but the condition is estimated to affect approximately 2-4% of the global population, with a three-times higher prevalence among women than men ^2,3^. FMS is considered a multifactorial condition influenced by genetic, environmental, psychological, and neurobiological factors resulting in central pain processing dysfunction ^4–8^. Central sensitisation amplifies nociceptive responses, lowering pain thresholds and exacerbating symptoms ^9^. Neuroimaging studies reveal structural brain changes, such as reduced grey matter in pain-modulation regions and increased density in other areas, alongside hyperactivation of key pain-processing areas like the somatosensory cortex ^10–14^. Diagnosis remains a challenge due to the absence of definitive biomarkers and is further complicated by symptom overlap with other conditions, resulting often in long diagnostic delays or misdiagnosis ^15^. Current treatment approaches focus on symptom management, such as pain relief, sleep improvement, and psychological support, rather than targeting the underlying disease mechanisms ^16–18^. These challenges underscore the urgent need for deeper understanding of the mechanisms driving FMS. Importantly, mechanism-based diagnostic tools, and treatments, will likely deliver a seismic change in these patients’ quality of life.

Growing evidence implicates the immune system in the pathophysiology of various neurological conditions, including those associated with pain. Immune cells such as B cells, T cells, and microglia interact with the central and peripheral nervous systems, influencing pain perception and modulation ^19^. Aberrant immune responses, characterised by altered cytokine profiles, chronic inflammation, and autoimmunity, have been linked to neurological conditions that include pain-sensitisation such as multiple sclerosis, chronic fatigue syndrome, autoantibody-driven encephalitis and neuropathic pain disorders.

FMS is increasingly recognised as involving chronic, low-level immune activation. Elevated levels of pro-inflammatory cytokines such as TNF-α, IL-6, and IL-8 are frequently observed ^20^ and correlate with symptom severity ^21–23^. Transcriptomic analyses have revealed heightened activation in T-helper and B cell pathways ^24,25^, with immune signatures resembling those seen in systemic autoimmune diseases. Interferons (IFNs), particularly IFN-γ, are consistently upregulated in FMS and may drive both neuronal sensitization and immune dysfunction. IFN-stimulated gene expression is increased in peripheral blood, resulting in disrupted tolerance and autoantibody production ^24,25^, mechanisms commonly implicated in autoimmune conditions like lupus and rheumatoid arthritis (RA) ^27,28^. Additionally, altered gut microbiome and bile acid profiles in FMS may further impact immune regulation ^29,30^, potentially via interferon signalling ^31^.

Emerging data also suggests that B cells may play a significant role in the pathogenesis of FMS ^32,33^. As key players in the adaptive immune system, B cells contribute to antibody production, antigen presentation, and cytokine secretion. Their surface-expressed B cell receptors (BCRs), which function as antibodies when secreted, are essential to immune recognition and regulation. These have been shown to exhibit abnormalities in FMS patients: IgG antibodies from FMS patients bind to human and murine dorsal root ganglia (DRG), and induce sensitivity to both painful and non-painful stimuli in mice^32^. Furthermore, elevated levels of anti-satellite glial cell IgG are linked to greater pain and Fibromyalgia Impact Questionnaire (FIQ) scores ^34^. Together, these findings support a model of FMS as a disorder of immune dysregulation, with overlapping features of autoimmunity.

Maintaining B cell tolerance is crucial to preventing autoimmunity. Peripheral mechanisms such as clonal deletion, receptor editing, and functional anergy eliminate or silence autoreactive B cells ^35–39^. Functional anergy represents a critical checkpoint where autoreactive B cells become unresponsive to antigen stimulation while remaining in circulation. Breakdown of these tolerance checkpoints is a hallmark of autoimmune diseases, enabling pathogenic B cell clones to produce autoantibodies and potentially drive chronic inflammation. Although B cell tolerance defects have been documented in other chronic pain-related autoimmune conditions, such as systemic lupus erythematosus (SLE) ^40^, no such studies have been reported in FMS to date.

In this paper, we intend to investigate this question, whether B cell tolerance in FMS is fundamentally compromised. By integrating high-resolution BCR repertoire sequencing, deep phenotyping, and functional assays in a well-characterised patient cohort, we uncover profound defects in peripheral B cell tolerance, including failed anergy and dysregulated BCR heavy chain clonal selection mirroring autoimmune pathology. These findings not only redefine FMS as a disorder of immune dysregulation but also unveil novel therapeutic opportunities to restore B cell homeostasis, offering hope for mechanism-driven interventions in this debilitating condition.

## Results

### Distinct class-switching in FMS

To investigate if B cells are dysregulated in FMS, we analysed the BCR repertoires of peripheral blood mononuclear cells (PBMCs) from 15 individuals with FMS and 19 healthy matched controls (**Supplementary Table 1**). Patients were required to meet the ACR 2010 or 1990 criteria for enrolment, and to have a pain intensity of at least 4/10 (11-point numeric rating scale with 0=no pain, 10=pain as bad as one can imagine); we only included patients with greater than 1 year disease duration. Patients had been assessed by a consultant in pain medicine and typically by a consultant rheumatologist and had been excluded if they suffered from established autoimmune conditions such as rheumatoid arthritis. Age and body mass index (BMI) were slightly higher in the FMS group, therefore, all subsequent statistical tests used age and BMI as a covariate. Co-morbid conditions of depression and mental health disorders, (non-inflammatory/non-rheumatoid arthritis) arthropathies and joint disorders, and gynaecological and obstetric disorders were significantly enriched in the FMS patients. This finding is in line with the established clinical literature on FMS co-occurrence^41^ and does not suggest a unique bias in our recruitment. Crucially, no healthy controls in either the BCR or FACS cohorts reported comorbidities or medications related to primary or secondary immune deficiencies or active autoimmune diseases. Likewise, the usage of analgesics, antidepressants, and gastrointestinal medications were significantly enriched in the FMS cohort. This pattern is also consistent with the typical management of FMS patients and does not introduce confounding immune deficits. Patients with FMS were majority female (82% and 89% in FMS and healthy control groups respectively) and had experienced widespread pain for 14 years (range 3-45 years) The patients’ *current pain* intensity at assessment was mean 7.47 (range 6-10), compared to 0 (SD=0) for the control group, and their mean (pressure pain threshold) PPT kPa over the lateral forearm was 176.087 (SD=64.776) compared to 468.421 (SD= 186.175) for the control group (p-values<1e-4).

BCR sequences encoding both the antigen-binding (VDJ) and constant regions of the BCR heavy chain were amplified ^42^ (**Figure 1a, Supplementary Table 1-2**, post-filtering BCR sequence capture range 3,293-84,298), facilitating analyses of isotype class and subclass, as well as allowing the quantitation of clone frequency and correction of PCR- or sequencing-based error. From this, metrics describing isotype usages, heavy chain clonality, level of somatic hypermutation (SHM) and features of the most variable regions of the BCR sequence (complementary determining region 3 (CDR3)) were calculated across all patients (**Figure 1b, Supplementary Figure 1**). Overall isotype usages were distinct between groups, with lower percentage of IGHG2 in FMS patients. When considering BCRs from antigen experienced isotype-switched B cells (i.e. from B cells expressing IGHA, IGHG or IGHE isotypes), a clear elevation of IGHA2 was additionally observed (**Figure 1b-c**). However, no detectable differences in class-switch recombination (CSR) was observed (through quantifying the frequency of unique VDJ regions that share two isotypes, suggesting their common clonal origin, after normalising for read depth ^42^, **Supplemental Figure 1l**), suggesting either historical or tissue-specific differences in the CSR process that were not detected in blood. Together this points towards differential class-switching in FMS skewed towards IGHA1 and IGHA2 rather than IGHG3.

**Figure 1.**
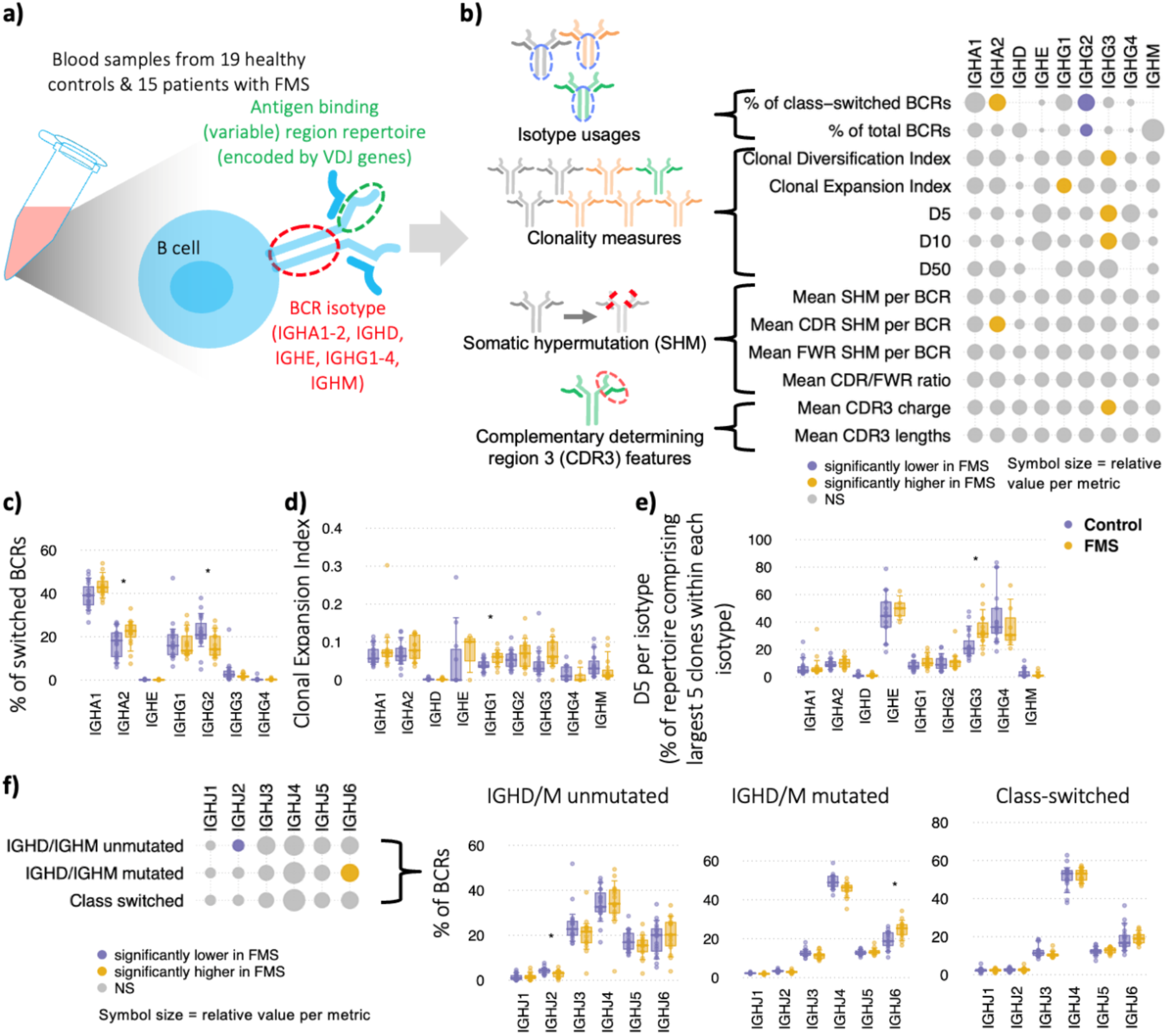
Altered B cell receptor (BCR) heavy chain selection and repertoire features in FMS. **a)** Overview of BCR sequencing workflow. RNA was extracted from peripheral blood mononuclear cells (PBMCs) from 15 individuals with fibromyalgia syndrome (FMS) and 19 age- and sex-matched healthy controls. BCR heavy chain transcripts spanning both the variable (VDJ) and constant regions were amplified. Repertoire metrics were derived, including isotype distribution, heavy chain clonal expansion, somatic hypermutation (SHM), and properties of the complementarity-determining region 3 (CDR3). **b)** Heatmap summarising differences in BCR repertoire features between FMS and healthy controls across isotypes. Yellow circles indicate significantly elevated frequencies in FMS; purple circles denote enrichment in controls. Circle size reflects the relative mean difference. **c-e)** Boxplots of selected BCR heavy chain features from panel b: **(c)** proportion of switched isotypes, **(d)** Clonal Expansion Index, and **(e)** D5 (diversity metric representing the fraction of dominant clones accounting for 5% of the repertoire). **f)** Heatmap of IGHJ gene usage frequencies, stratified by unmutated IGHM/D (representing predominantly naïve B cells), and mutated IGHM/D and class-switched BCRs (representing antigen experienced B cells). (Right) Boxplots of these distributions, with accompanying boxplots showing distributions across groups. Statistical testing in b-d used two-sided MANOVA; * denotes p < 0.05. Boxplots represent the 10th, 25th, 50th (median), 75th, and 90th percentiles. FMS: n = 15; Healthy Controls: n = 19.

### Distinct heavy chain clonality and SHM in FMS

B cell clones are defined by sharing a unique VDJ rearrangement and can be characterised by size (clonal expansion) and diversification (owing to SHM and isotype switching). Both the clonal diversification index (a measure of the unevenness of unique VDJ region sequences per clone ^42^) and the clonal expansion index (a measure of the ‘unevenness’ of the number of RNA molecules per unique VDJ region sequence ^42^) were significantly elevated in FMS patients in IGHG3 and IGHG1 isotypes, respectively (**Figure 1b, d)**. Other clonality metrics measuring the proportion of the repertoire accounting for the top 5 (D5) and top 10 clones (D10) per isotype demonstrate that polyclonal expansions are elevated in FMS in IGHG3, an effect not observed in the top 50 clones (D50) (**Figure 1b, e**). Together this indicates a broad increase in IGHG B cell clonality in FMS patients rather than selected dominant clones.

### B cell selection defects in FMS resemble autoimmune disorders

Given the differences in isotype usages and heavy chain clonality evidenced in antigen-experienced (non-naïve, memory) B cells, we next investigated if B cell selection was distinct between groups. Given that different IGHV and IGHJ genes have a propensity to bind different groups of antigens, alternative IGHV and IGHJ usages are indicative of differences in B cell selection. Unlike in classical autoimmune diseases ^42^, there were no significantly different IGHV usages in BCRs from naïve B cells (i.e. IGHD/M with no SHM (unmutated), **Supplemental Figure 2a**). However, there were some elevated IGHV gene usages observed in antigen-experienced B cells (i.e. from class-switched BCRs), including IGHV6-1 in IGHA1, IGHG1 and IGHG3 isotypes, an observation also previously seen in other autoimmune diseases including systemic lupus erythematosus (SLE), Crohn’s disease and eosinophilic granulomatosis with polyangiitis (EGPA) ^42,43^. Interestingly, elevated levels of IGHV4-34 proportions in IGHG3 BCRs were observed (**Supplemental Figure 2a-b**), which is known to have elevated autoreactivity and polyreactivity in multiple autoimmune diseases driven by framework region 1 (FR1) AVY and complementary determining region 2 (CDR2) NHS amino acid motifs ^42,44^.

Next, we explored whether IGHV- and IGHJ-specific clonal expansion differed in FMS. Clonal expansion of the specific IGHV genes (measured by mean clone percentage of repertoire) was found to be significantly higher in FMS patients compared to healthy controls, most notably in the class-switched B cells (**Supplemental Figure 2c**). This effect was most pronounced in the IGHD/M mutated BCRs and class-switched BCRs with 9 and 8 of the IGHV genes, respectively, showing significant differences between patient groups. In contrast, 4 IGHV genes showed significant differences in the expanded clones (defined as >1% of total repertoire). Interestingly, the clonal expansion of class-switched IGHV4-34 expressing B cells was significantly higher in FMS patients compared to controls. On the other hand, for BCRs containing genes seen at a lower frequency in FMS patients, namely IGHV1-18, these were found to also have a higher level of clonal expansion in the class-switched B cells. Finally, the IGHV-specific levels of somatic hypermutation were also found to be significantly elevated between FMS and healthy controls, most notably in IGHV4-59 BCRs (for IGHA2 and IGHG3) and IGHV1-18 and IGHV1-69 (for IGHG3) (**Supplemental Figure 3a-b**). Taken together, these data strongly suggest that FMS patient B cells have an altered, IGHV-specific propensity for clonal expansion and SHM upon activation and class-switching, indicating a differential response of specific B cell subsets to chronic stimuli in FMS. Finally, the IGHJ gene usage showed strongly significant distinctions between groups, with FMS patients displaying elevated IGHJ6 gene usage (**Figure 1d, Supplemental Figure 3c)**. Interestingly, this is observed only in the antigen-experienced IGHD/M B cells (i.e. that have undergone SHM), suggesting differences in peripheral tolerance. Indeed, a reanalysis of the BCR repertoires from Bashford-Rogers *et al.* ^42^ revealed elevated usage of IGHJ6 in other autoimmune diseases including anti-neutrophil cytoplasmic antibody (ANCA)-associated vasculitis (AAV), Crohn’s disease and SLE, with the strongest effect observed in antigen-experienced IGHD/M mutated B cells (**Supplemental Figure 4**). Finally, alterations in the complementarity-determining region 3 (CDR3) of the BCR, including increased length and positive charge, have been associated with antibody polyreactivity and autoimmunity ^45^. Consistent with patterns observed in other autoimmune conditions, mean CDR3 charge was elevated in FMS compared to controls. However, in contrast to these conditions, CDR3 lengths were not significantly different in FMS (**Figure 1b**).

Overall, these findings reveal that antigen-experienced B cells in FMS patients exhibit altered selection patterns, characterised by differential usage of selected IGHJ and IGHV genes, including IGHJ6 and the autoreactive IGHV4-34, a feature shared with other autoimmune diseases. IGHV-specific clonal expansion and SHM was observed. Notably, while significant differences were detected in antigen-experienced B cell selection, clonality, and isotype switching, no such differences were observed in the naïve B cell compartment (IGHD/M BCRs). This suggests that FMS patients primarily display alterations in peripheral B cell selection.

### Elevated naïve B cell proportions in FMS patients

Given that the BCR repertoire in FMS patients shows evidence of altered peripheral tolerance, manifesting as changes in BCR-specific activation, class switching, and clonal expansion, we next investigated whether the composition of B cell populations and their responses to stimulation were also affected. From a subset of patients from the BCR cohort FMS: n=14, healthy controls: n=14, with age and BMI not significantly different between groups (T-test, **Supplementary Table 1**), PBMCs were thawed and subjected either to flow cytometry analysis immediately (day 0, representing cellular populations *within donors*), or to a 5-day *ex vivo* stimulation with subsequent flow cytometry analysis including proliferation measurements (day 5, **Figure 2a**). To assess B cell responsiveness without introducing BCR-dependent bias, we used CpG + IL-2 stimulation, a potent BCR-independent activator of naïve human B cells^46–48^. This avoided the methodological confounding inherent to anti-IgM stimulation, which disproportionately activates IgM⁺⁺ cells and would obscure true anergy-associated signalling defects. CpG (TLR9 agonist) triggers strong proliferation and differentiation independently of surface IgM abundance, thereby providing an unbiased measure of intrinsic activation capacity across different IgM levels subsets. In addition, stimulating the full PBMC mixture rather than purified B cells enabled us to model a physiologically relevant, multi-cellular activation environment. CpG activates antigen-presenting cells to upregulate CD40 and produce cytokines (e.g., IL-12, type I IFN), generating a robust T-dependent–like co-stimulatory milieu that more closely reflects autoimmune-associated immune activation ^49–53^. This approach allowed us to detect subtle differences in B-cell activation and tolerance mechanisms that may only emerge under complex co-stimulation, thereby providing a more translationally meaningful assessment of B-cell dysregulation in FMS.

**Figure 2.**
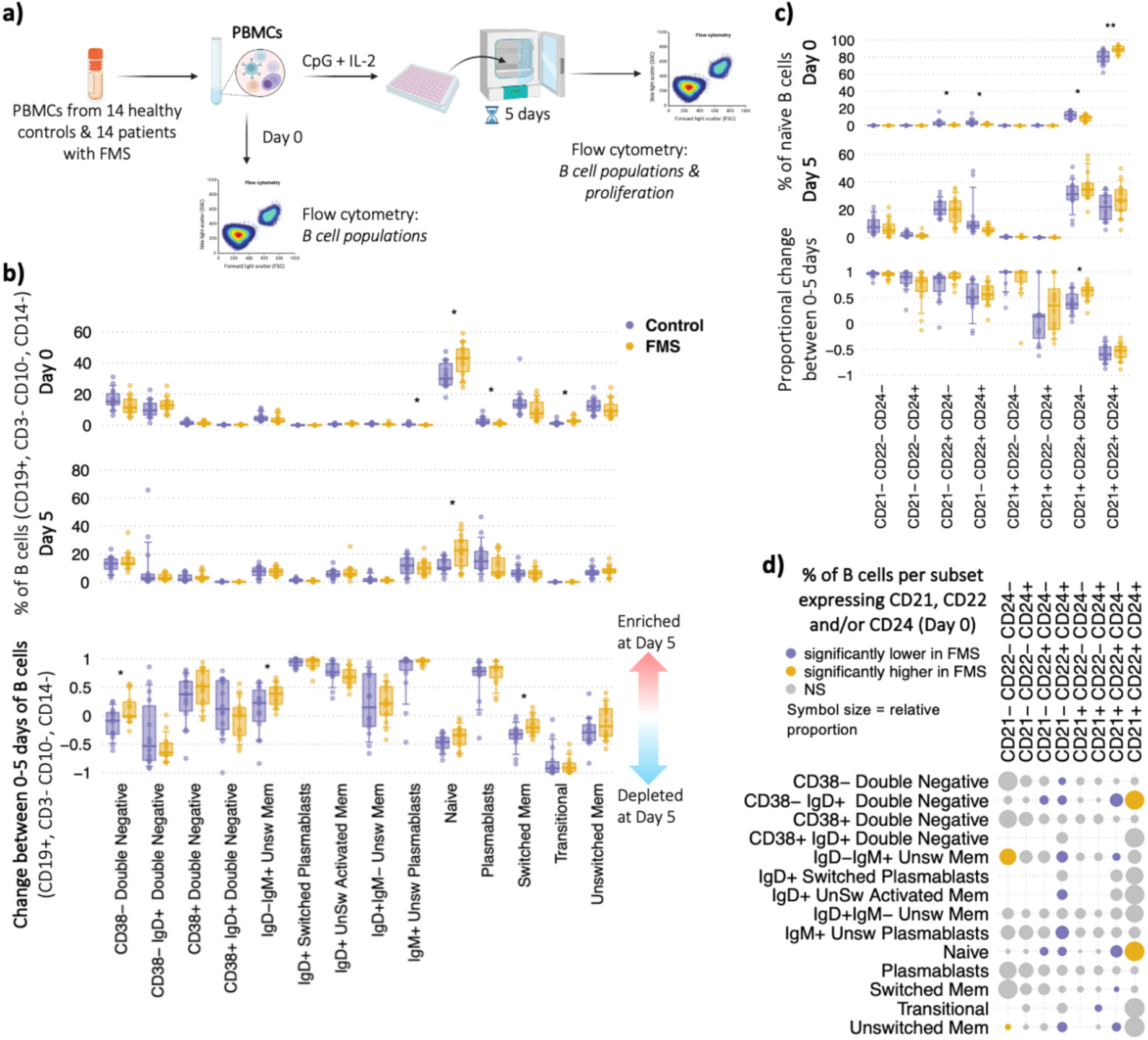
Altered B cell fate and surface phenotype in FMS. **a)** Experimental workflow. Peripheral blood mononuclear cells (PBMCs) from fibromyalgia syndrome (FMS) patients (n = 14) and healthy controls (n = 14) were split into two arms: immediate flow cytometry profiling (day 0) and *ex vivo* stimulation for 5 days followed by flow cytometry (day 5). **b)** Frequencies of canonical B cell subsets within the CD19⁺ population across groups at day 0 and day 5, and their respective proportional change between timepoints. **c)** Distribution of CD21/CD22/CD24 combinations within naïve B cells at day 0 and day 5 in FMS versus controls, and their respective proportional change between timepoints. **d)** Heatmap summarising differences in the distribution of CD21, CD22, and CD24 surface marker combinations across B cell subsets. Yellow circles indicate significantly elevated frequencies in FMS; purple circles denote enrichment in controls. Circle size reflects the relative mean difference. Statistical testing used two-sided MANOVA; * denotes p < 0.05, ** denotes p < 0.005. Boxplots represent the 10th, 25th, 50th (median), 75th, and 90th percentiles. FMS: n = 14; Healthy Controls: n = 14.

B cells were gated on CD19+, CD3- CD10-, CD14-, and phenotypes based on canonical B cell subsets ^54^ (gating in **Supplemental Figure 5, Supplementary Table 3**). Whilst no differences were seen in the proportion of live cells that were B cells between FMS patients and controls (**Supplemental Figure 6a**), we observed significant differences in the proportions of B cell subsets (**Figure 2b**). Most notably, FMS patients had a significantly higher proportion of naïve and transitional B cells at day 0 (p-value=0.0113 and 0.0123 respectively) than healthy controls, and significantly lower proportions of plasmablasts and IgM+ Unswitched plasmablasts (p-values<0.03). Indeed, naïve B cells were also significantly elevated in FMS patients at day 5 post-stimulation (p-value=0.0103, **Figure 2b**). The increased levels of naïve B cells observed in FMS patients align with previously reported tolerance checkpoint defects that shape the naïve B cell repertoire in other autoimmune diseases ^35–38^. However, the relative changes in B cell subset proportions between day 0 (pre-stimulation) and day 5 (post-stimulation) were comparable between FMS patients and controls (**Figure 2b**), indicating no detectable differences in B cell differentiation.

### Elevated CD21, CD22 and CD24 expression in naïve B cells in FMS

Peripheral B cell tolerance, a critical safeguard against autoimmunity, is primarily maintained through mechanisms such as anergy and apoptosis^55^. Anergic B cells are characterised by a state of unresponsiveness to antigenic stimulation, exhibiting dampened BCR signalling and reduced effector functions, reduced proliferative capacity, lower cytokine and antibody secretion, and a shorter lifespan^5^, often resulting from prolonged BCR engagement with self-antigens. Defects in anergy mechanisms are a frequent hallmark of autoimmune pathology^57^. Molecular regulators implicated in maintaining anergy include CD22, FCGR2B, and TNFAIP3, all of which exert inhibitory control over BCR signalling and whose reduced function or expression associates with autoimmune disease in both humans and mice. Functionally, anergic B cells display altered tissue localisation, and impaired T cell interactions^56,58–62^. CD21 (Complement Receptor 2) amplifies BCR signalling, making its downregulation a consistent marker of lower responsiveness (anergy)^63^. CD21 downregulation has been consistently linked to lower responsiveness to stimulation, and observations of enriched populations of CD21^low^ naïve B cells have been shown in autoimmune diseases such as rheumatoid arthritis^64^. CD22 serves as a negative regulator of BCR signalling, crucial for maintaining anergy and immune tolerance, with its absence leading to B cell hyperresponsiveness^65^. CD24 similarly functions as an inhibitory regulator, dampening B cell activation^66–68^.

Defects in CD21, CD22 and CD4 expression have been shown in autoimmune diseases^37,64^. To determine if similar tolerance defects are present in FMS, we systematically assessed the expression of CD21, CD22 and CD4 across the B cell populations. We found that while over 60% of naïve B cells in both groups expressed this CD21+CD22+CD24+ phenotype, FMS patients exhibited a significantly higher proportion of these cells compared to healthy controls (**Figure 2c-d**). This increased baseline co-expression is highly suggestive of fundamental alterations in peripheral tolerance mechanisms in FMS. The elevated presence of B cells poised for activation, expressing co-receptors that promote CD21 or dampen CD22/CD24 B cell signalling, indicates a compartment with a lowered threshold for activation. Furthermore, upon *in vitro* BCR stimulation, CD24 expression was downregulated in naïve B cells, as anticipated^37^. Critically, FMS patients displayed a significantly higher increase of CD24-naïve B cells post-stimulation (**Figure 2c**). The loss of this negative regulatory marker upon stimulation strongly implies an increased propensity for activation and hyperresponsiveness in FMS naïve B cells, reinforcing the notion of a dysregulated compartment. Significant differences in CD21, CD22 and CD24 expression were also observed in antigen-experience B cell subsets including CD38-IgM?+ double negative (CD27-IgD-) B cells, IgD-IgM+ unswitched memory B cells, and unswitched memory B cells (**Figure 2d, Supplemental Figures 6b-d**). These findings suggest that naïve B cells in FMS patients may be dysregulated, leading to changes in differentiation and proliferative potential.

### Defective functional anergy in FMS patients

Anergic naïve B cells have been defined primarily by low surface IgM (IgM^lo^) expression, a feature first characterised in murine models^69^ and later confirmed in humans ^56,57,70,71^. These cells have been shown to display impaired BCR signalling upon IgM or IgD engagement, reduced activation and costimulatory molecule expression (CD19, CD21), increased inhibitory CD22 levels, and diminished proliferative and differentiation capacity^56^. IgM^lo^ naïve B cells have been shown to exhibit distinct repertoire features, including longer CDR3 regions and increased IGHJ6 usage, signatures linked to polyreactivity and autoimmune potential^45,57^, similar to the defects observed in the BCR repertoires sequencing data in FMS (**Figure 1**). Collectively, these studies suggest that IgM^lo^ naïve B cells represent a restrained naïve B cell compartment critical for maintaining peripheral tolerance requiring the influence of co-receptors CD21, CD22 and CD24. To specifically assess functional naïve B cell anergy in FMS, we first confirmed that low surface IgM (IgM^lo^) expression serves as a robust *in vivo* surrogate marker for the anergic phenotype in healthy individuals^56,57,69,70^. Our re-analysis of single-cell multi-omic data from 8 healthy controls, comprising 1,192 high-confidence naïve B cells (from COMBAT^72^, mean cells per patient: 301.2, range: 139-466, **Figure 3a-b**) from which we calculated the proliferation score and BCR signalling score based on GOBP_B_CELL_PROLIFERATION and KEGG_B_CELL_RECEPTOR_SIGNALING_PATHWAY gene sets respectively. These give a read out of the proliferative and BCR signalling nature of these cells based on the expression of genes within these gene sets. Across all the top 29 highest expressed protein markers in naïve B cells (out of 192 proteins), we found that the strongest correlates of BCR signalling were IgD, CD1c, CD24, and IgM (p-values<1e-5, **Figure 3c, Supplementary Figure 7a-b**), in agreement with previous studies. Other positively associated markers were CD45RA and β7-integrin, both of which have also been shown to have roles in B cell tolerance^73,74^. Indeed, the B cells with <33^rd^ percentile BCR signalling and proliferation scores exhibited lowest IgM expression compared to the B cells in the top 33^rd^ percentiles for each score (**Figure 3d-e, Supplementary Figure 7c**). BCR signalling was significantly correlated with B cell proliferation signatures (p-value= 1.31E-27, **Figure 3c**), and CD22, CD19, CD24, FCGR2B, CR2 (CD21) transcripts. Together, these results provide functional confirmation that low IgM is a reproducible surrogate for higher levels of anergy in naïve B cells *in vivo*, which correlates with lower CD21, CD22, and CD24 as well as BCR signalling and proliferation.

**Figure 3.**
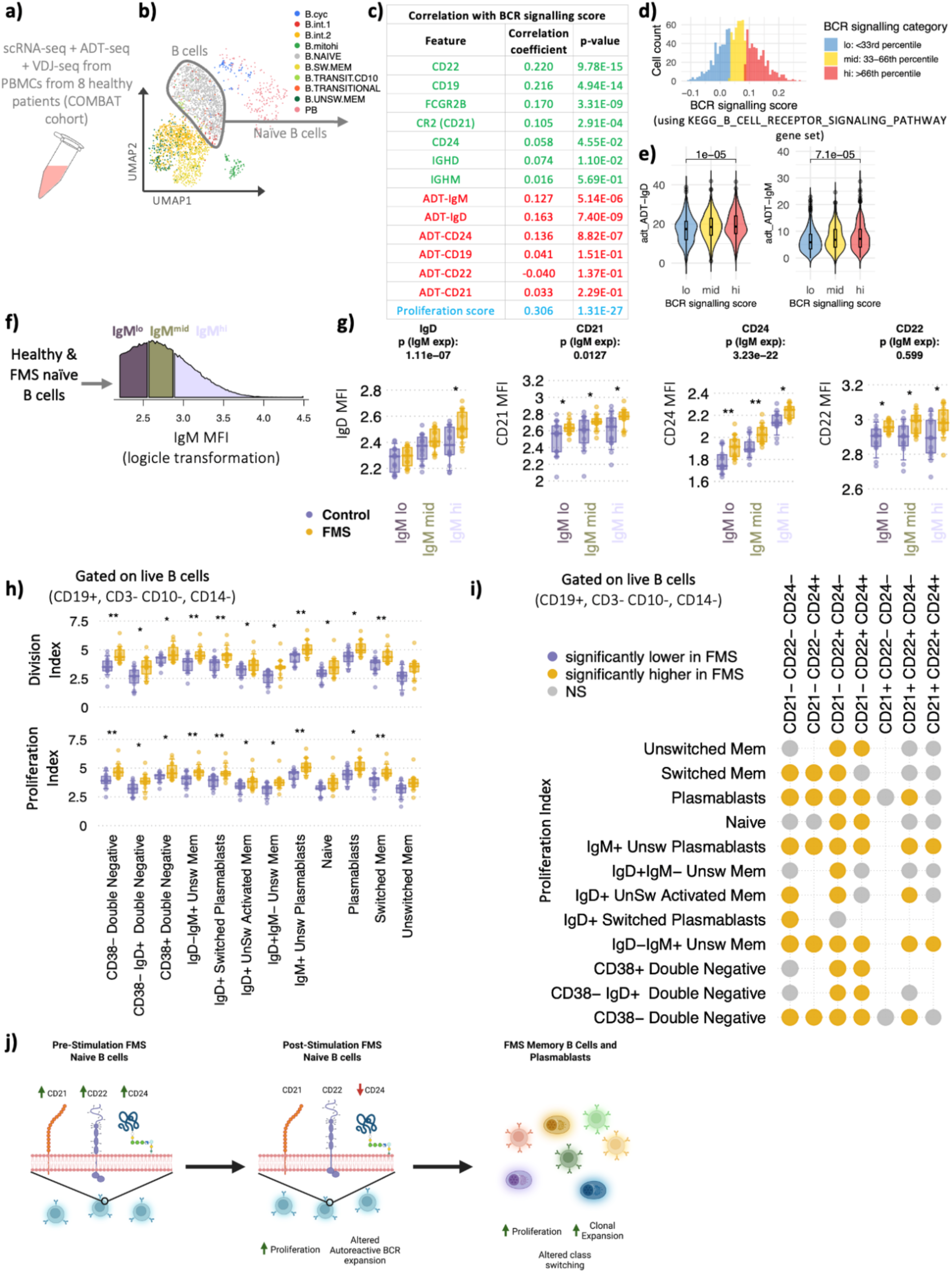
Impaired functional anergy and enhanced B cell proliferation in FMS. Schematic showing the **a)** re-analysis of single-cell multi-omic data from COMBAT using 8 healthy controls, **b)** comprising 1,192 high-confidence naïve B cells. **c)** Correlation of cell-surface protein expression (green), gene expression (red) and proliferation score (blue) with BCR signalling score. **d)** Naïve B cells were divided into the lowest, mid and highest BCR signalling score thirds and **e)** correlated these cell populations with surface IgD and IgM expression via cell violin plots. **f)** FMS and healthy control B cells from flow cytometry were divided into IgM-lo, -mid, -hi naïve B cells. **g)** Surface expression levels of IgD, CD21, CD22, CD24 of FMS and healthy controls within IgM-lo, -mid, -hi naïve B cells, grouped by patient group. P-values at the top of each plot gives the overall correlation of each marker with IgM expression. **h)** Boxplots of selected proliferation indices per B cell subset groups at day 0 and day 5, and their respective proportional change between timepoints. **i)** Heatmap summarising differences in the proliferation index after 5 days of *ex vivo* stimulation, stratified by CD21, CD22, and CD24 surface marker across B cell subsets. Yellow circles indicate significantly elevated frequencies in FMS; purple circles denote enrichment in controls. Circle size reflects the relative mean difference. **f)** Summary schematic illustrating key findings of altered anergy and proliferation dynamics in FMS B cells. Created in BioRender. Wu, S. (2025) https://BioRender.com/fyyca2i. Statistical testing used two-sided MANOVA; * denotes p < 0.05, ** denotes p < 0.005. Boxplots represent the 10th, 25th, 50th (median), 75th, and 90th percentiles. FMS: n = 14; Healthy Controls: n = 14.

Acknowledging that naïve B cell anergy should be considered as a continuum rather than a set of discrete cellular states, we compared the phenotypic extremes of this spectrum in FMS and healthy controls. The naïve B cells were stratified based on IgM expression levels (IgM^lo^ with <33^rd^ percentile IgM expression, IgM^mid^ with 33^rd^-66^th^ percentile IgM expression and IgM^hi^ with >66^th^ percentile IgM expression (**Figure 3f**). While the overall proportion of IgM^lo^ cells was unchanged between patient groups (**Supplemental Figures 7d-e**), suggesting the *number* of anergic cells is not reduced, we observed a profound functional defect. When accounting for the level of IgM expression, FMS patients exhibited significantly elevated expression of CD21, CD22, and CD24 across all IgM expression levels (**Figure 3g**). Most strikingly, the most anergic IgM^lo^ naïve B cells in FMS patients exhibited the most significantly elevated expression of CD21, CD22, and CD24. This is a critical finding: the B cells that are supposed to be the most anergic (defined by IgM^lo^) display markers that should be low for this phenotype (CD21^lo^) or are part of the inhibitory machinery (CD22 and CD24). No differences between FMS and healthy controls were observed after 5 days of stimulation (**Supplemental Figures 7f**). This suggests a failure to adequately execute the anergic program in FMS B cells across IgM expression levels. The sustained high expression of these key receptors on the most anergic population (IgM^lo^ B cells) indicates a diminished functional anergy and an increased, unrestrained responsiveness to stimulation in FMS, representing a clear breakdown in peripheral B cell tolerance.

### FMS patients have more proliferative B cells

Finally, to test whether B cells from FMS patients are less anergic and more responsive to stimulation, we examined B cell proliferation following 5 days of stimulation. FMS patients displayed significantly increased proliferation across most B cell subsets, as quantified by both proliferation and division indices (**Figure 3h-i, Supplemental Figures 8**). All CD24-B cell subsets, as well as IgD⁻IgM⁺ unswitched memory B cells, showed increased proliferation following stimulation. Notably, these effects were most pronounced within CD21⁻ and CD24⁻ B cell populations, reinforcing a connection between altered CD21/CD24 expression and heightened activation potential. Similar patterns of excessive B cell activation have been observed in systemic autoimmune diseases, where defective tolerance mechanisms drive inappropriate immune responses ^75^. The increased proliferation observed in FMS further supports a model in which anergic B cell dysregulation contributes to autoantibody pathology, aligning with broader autoimmune paradigms.

Overall, we have demonstrated defective anergy in FMS B cells, with elevated naïve B cell proportions, heightened CD21, CD22, and CD24 expression, even when accounting for the same IgM expression suggesting compromised tolerance checkpoints contribute to FMS pathogenesis (**Figure 3j**). Upon stimulation, the naïve B cells more readily reduced their CD24 expression, a known negative regulator of B cell activation ^68^, and exhibited exaggerated proliferative responses. We observed a selective expansion of B cells expressing BCRs linked with autoreactivity and significantly higher proliferation. Memory B cells and Plasmablasts were also more proliferative, with evidence of dysregulated clonal expansions, class-switching (skewed toward IGHA1) and somatic hypermutation, and altered selection patterns (increased IGHV6-1/IGHJ6 usage) resembling autoimmune diseases ^42,43^. Our findings reveal a pattern of B cell dysregulation in FMS that mirrors key features of autoimmunity. The combination of defective anergy, heightened activation potential, and aberrant clonal expansion suggests a breakdown in immune tolerance, potentially contributing to disease pathology. The shared genetic architecture between FMS and immune-related pathways further supports an immunological basis for FMS, with FMS risk-associated genes linked to immune cell activation, microbiome interactions, and interferon responses (**Tables S4-5**) derived from the GWAS catalog ^76^. These results highlight a critical role for B cell dysfunction in FMS and suggest potential avenues for targeted immunomodulatory interventions.

## Discussion

The findings of this study provide compelling evidence that FMS is characterised by profound defects in peripheral B cell tolerance, mirroring immunological dysregulation observed in classical autoimmune diseases. Our integrated analysis of BCR repertoires, phenotypic profiling, and functional assays reveals three key perturbations: (1) skewed class-switching toward IGHA2 and not IGHG3; (2) elevated clonal expansion and diversification, (3) altered selection patterns (increased IGHV6-1/IGHJ6 usage, differences in IGHV-specific clonal expansion, differences in SHM) in antigen-experienced B cells, (4) defective naïve B cell anergy marked by elevated CD21/CD22/CD24 expression and hyperproliferation, and (5) polyclonal expansion in class-switched B cells. These results fundamentally redefine FMS as a disorder of immune tolerance, challenging its historical classification as purely a central pain sensitisation syndrome.

The observed shifts in isotype usage, particularly the IGHA1 skewing and reduced IGHG2, parallel patterns seen in systemic lupus erythematosus (SLE) and rheumatoid arthritis (RA), where these aberrant class-switching drives pathogenic autoantibody production ^42,77^. Notably, the elevated IGHJ6 usage in mutated IGHD/M+ B cells aligns with defects in early peripheral tolerance, as reported in ANCA-associated vasculitis ^42^. While FMS lacks the overt autoreactive CDR3 features (e.g., elongated CDR3) of diseases like SLE, the convergence of clonal expansion, SHM dysregulation, and previously published interferon signatures ^26^ suggests a shared mechanism of B cell checkpoint failure. This is further supported by the increased naïve B cell compartment, a feature linked to defective central tolerance in autoimmune-prone individuals ^35–38^.

Critically, the phenotypic and functional data indicate a disruption in B cell anergy mechanisms in FMS. Naïve B cells from FMS patients exhibit elevated expression of CD21, CD22, and CD24, markers associated with BCR signalling regulation, even after accounting for IgM levels. This contrasts with the CD21^lo^ anergic phenotype typically observed in healthy individuals, which is associated with reduced BCR responsiveness and immune tolerance of autoreactive clones ^64^. Elevated CD21 expression, which enhances BCR signalling, in the context of lower surface IgM, suggests these cells are unusually primed for activation. Consistently, they exhibit exaggerated proliferative responses upon stimulation.

However, the implications of concurrently increased expression of CD22 and CD24, both negative regulators of BCR signalling ^68^, are less straightforward. As suggested in prior studies, the balance between activating and inhibitory signals is key to determining the maintenance of anergy or its breakdown. In this context, our observation of increased CD24 downregulation upon stimulation in FMS may reflect a shift in this balance toward activation, enabling hyperproliferation despite elevated CD24 levels at baseline. This underscores the idea that anergy is not simply defined by static marker expression, but by the dynamic interplay between activation and inhibition thresholds. Notably, this dysregulated phenotype parallels preclinical autoimmune states such as rheumatoid arthritis, where self-reactive B cells escape tolerance checkpoints despite inhibitory cues ^39^. At present, it is not fully established whether the increased expression of CD21, CD22, and CD24 on naïve B cells in FMS is driven intrinsically by the B cells themselves or by extrinsic soluble factors. Intrinsic factors may include genetic or epigenetic programming of B cells that alters baseline expression of key co-receptors, as observed in other autoimmune diseases where variants in the gene encoding CD21 (CR2) or CD22 influence expression and signalling. In parallel, extrinsic factors such as inflammatory cytokines, type I interferons, and other soluble mediators present in the serum of patients can rapidly modulate co-receptor expression on otherwise healthy B cells. FMS is associated with elevated levels of pro-inflammatory cytokines (IL-6, TNF-alpha) and chemokines^78^. These soluble factors can directly bind to receptors on B cells and signal changes in co-receptor expression^79^.

These findings bridge longstanding gaps in FMS research. The dysregulated BCR repertoires and hyperproliferation may explain reports of FMS IgG inducing pain hypersensitivity in mice ^32^. The IGHA1 skewing could reflect mucosal immune triggers, potentially linking to gut dysbiosis ^80,81^. The interferon signatures observed in previous studies may perpetuate B cell dysfunction via TLR7/9 activation ^26^, creating a feedforward loop of immune stimulation. While not yet diagnostic, the BCR repertoire PCA separation highlights immune profiling as a potential future tool to complement clinical criteria.

The overlap with autoimmune mechanisms suggests repurposing B cell-targeted therapies could be a viable option for some patients, such as anti-CD20 depletion of hyperactive naïve and memory B cells which may reset tolerance, as trialled in RA ^82^. Elevated BAFF (linked to interferon) promotes anergy escape, where BAFF inhibition through belimumab has been shown to improve fatigue in SLE ^83^, a key FMS symptom. Given interferon signalling’s role, JAK1/2 inhibitors (e.g., baricitinib) could dampen B cell activation ^84^.

While transformative, this study has caveats. Blood analyses may not capture tissue-resident B cell defects, particularly in gut or near affected nerves. Disease-driving auto-antigens have been so far elusive; however, the advent of newer high-throughput approaches will aid this work. Notably, there are still gaps in our understanding of why FMS, despite signs of immune dysregulation, does not exhibit the tissue-destructive autoimmunity typical of classic autoimmune diseases. Future studies leveraging longitudinal sampling will be critical to determine whether B cell dysregulation precedes symptom onset or disease flares. Additionally, stratifying larger patient cohorts by immune profiles may uncover distinct immunological endotypes. Exploring alternative immune stimulation strategies could further illuminate dysregulated pathways and intercellular communication contributing to FMS pathogenesis.

This study positions FMS within the spectrum of immune-mediated disorders, with defective B cell tolerance as a central driver. By identifying convergent autoimmune pathways, from interferon activation to anergy failure, we provide a roadmap for mechanistically grounded therapies, moving beyond symptomatic management toward disease modification.

## Materials and Methods

### Recruitment of study cohort

Diagnosed 15 well-phenotyped FMS patients and 19 healthy age-matched pain-free controls were recruited at The Walton Centre, UK as part of the Autoimmunity-informed Phenotyping in patients with FMS (APIF) study ISRCTN18414398. Study details have been published (PMID: 35652761). This study has received ethics approval from the Health and Care Research Board Wales (HCRW): 18/WA/0234, LBIH Biobank project number 18-10, approval date 5/10/2018). FMS patients were diagnosed based on American College of Rheumatology 1990 or 2010 diagnostic criteria or both. Samples were collected via venepuncture into EDTA blood bottles. PBMCs were extracted from blood samples of patients using a protocol adapted from the manufacturer ^85^. Extracted PBMCs were stored in liquid nitrogen. PBMC counts were approximately 1×10^7^ lymphocytes/mL per tube. Blood sample collection and PBMC extraction were performed by collaborators in University of Liverpool.

### Library preparation for BCR repertoire sequencing

PBMCs from patients were stored in liquid nitrogen prior to RNA isolation. PBMCs were thawed in a 37 °C bath for one minute, the diluted into 10ml pre-warmed Gibco™ RPMI cell culture medium via dropwise addition of cells. The cell suspension is centrifuged at 300 x g for 5 minutes to obtain cell pellet. Beta-mercaptoethanol was mixed with RLT buffer prior to RNA isolation. RNA isolation was conducted using the Qiagen® RNeasy Plus Mini kit according to the manufacturer’s protocol. The mRNA was then quantified using Qubit™ 4 fluorometer and stored in a -80 °C refrigerator.

BCRs were amplified using a protocol we have previously described ^42^. Briefly, RNA was reverse transcribed to cDNA using a mixture of IgA/D/E/G/M isotype specific primers, incorporating 15 nucleotide UMIs. The resulting cDNA was used as template for PCR amplification using a set of 6 FR1-specific forward primers including sample-specific barcode sequences (7 nucleotides) along with a reverse primer specific to the RT primer. Dual-indexed sequencing adapters (KAPA) were ligated onto ≤500 ng of amplicon per sample using the HyperPrep library construction kit (KAPA). The adapter-ligated libraries were finally PCR-amplified (initial denaturation at 95°C for 1 min, for 2-83 cycles at (98°C for 15 s, 60°C for 30 s, 72°C for 30s, and final extension at 72°C for 1 min). The libraries were sequenced on an Illumina MiSeq using the 2 x 300 bp chemistry.

The PCR amplicon pools were purified using AMPure beads, starting with a large sized purification step with 0.55x beads, followed by a small-sized purification step with 0.15x beads. Bead ratios were determined based on guidance from Beckman Coulter, Inc. and experimental optimisation. A washing step using 200μl 70% ethanol was performed, and DNA was then eluted in 55μl water. Gel electrophoresis was conducted on purified products to assess the efficiency of the clean-up step and to assess the quantity of the band of interest relative to the ladder. The library preparation is performed in accordance with the KAPA Hyper Prep Kit protocol. Based on Qubit™ quantification of libraries, all libraries were adjusted to the concentration of the library with lowest concentration. 5μl of each adjusted library was then pooled into a 50μl library pool with a final concentration of 62.73nM. Sequencing of the BCR library was performed by Azenta Life Sciences. One lane of 300bp paired end MiSeq was run in MiSeq2×300 flow cell with 15% PhiX Control v3 Library.

### BCR repertoire analysis

Raw sequencing reads were filtered for base quality (median Phred score >32) using QUASR^86^. Forward and reverse reads were merged if they contained an identical overlapping region of >50bp, or otherwise discarded. Universal barcoded regions were identified in reads and orientated to read from V-primer to constant region primer. The barcoded region within each primer was identified and checked for conserved bases. Primers and constant regions were trimmed from each sequence, and sequences were retained only if there was >80% per base sequence similarity between all sequences obtained with the same barcode, otherwise discarded. The constant region allele with highest sequence similarity was identified by 10-mer matching to the reference constant region genes from the IMGT database^87^, and sequences were trimmed to give only the region of the sequence corresponding to the variable (VDJ) regions. Isotype usage information for each BCR was retained throughout the analysis hereafter. Sequences without complete reading frames and non-immunoglobulin sequences were removed and only reads with significant similarity to reference IGHV and J genes from the IMGT database using BLAST^88^ were retained. Ig gene usages and sequence annotation were performed in IMGT V-QUEST, where repertoire differences were performed by custom scripts in Python.

### BCR isotype frequencies, somatic hypermutation, CDR3 lengths and IGHV/J gene usages

Analysis methods are based on ^42^. To account for the greater numbers of BCR RNA molecules per plasmablast compared to other B cell subsets, the we normalised isotype usages, defined as the percentage unique VDJ sequences per isotype, thus controlling for differential RNA per cell and reducing potential biases from differential RNA per cell. Similarly, mean somatic hypermutation levels and CDR3 lengths were calculated per unique VDJ region sequence to reduce potential biases from differential RNA per cell. IGHV gene usages were determined using IMGT, and proportions were calculated per unique VDJ region sequence. The representation of IGHV/J genes in the BCR repertoire reflects their presence in the germline, the naïve repertoire and their expansion after antigenic exposure. We therefore compared the frequency of IGHV gene use in PBMC-derived BCRs identified by sequence as being enriched for naive (IgM+D+SHM-: >78% naïve B cells by flow cytometry) and antigen-experienced B cells (including both unswitched (IgM+D+SHM+) and class-switched memory (IgA+/G+/E+) subsets) as shown in ^42^.

### PBMC isolation and culture

Cryopreserved PBMCs were removed from storage and immediately transferred to a 37°C water bath. Vials were held in the water bath without submerging the cap and gently agitated for 1 minute until a small ice crystal remained. Cells were then transferred to 15ml Falcon tubes with 10µl of DNAse I Solution (STEMCELL) and immediately diluted with prewarmed RPMI Medium 1640 (ThermoFisher) and centrifuged at 500 x g for 5 minutes. Once complete, the supernatant was removed, and Complete media (RPMI Medium 1640 + 10% Heat Inactivated Foetal Bovine Serum/FBS (ThermoFisher) + 1% Penicillin/Streptomycin (Thermofisher)) was added to the leftover pellet. Cells were then counted and either prepared for Day 0 staining or stained with eBioscience Cell Proliferation Dye eFluor 450 (Thermofisher) according to the manufacturer’s protocol. Post-proliferation staining, cells were counted again and the cell concentration was made to 2×10^6^ cells/ml for the seeding of 100µl of cell mix (i.e. 2×10^5^ cells) with 100µl of CpG ODN 2006 (Invivogen) and IL-2 (Biolegend) that reach a final concentration of 0.35µM and 100 IU/ml respectively. The cells were then left for 115-120 hours (∼5 days) in an incubator at 37°C and 5% CO_2_. After incubation, cells were harvested, counted and stained for flow cytometry with any excess PBMCs available being frozen in freezing media (45% FBS, 45% Complete Media & 10% DMSO).

### Flow cytometry staining and acquisition

PBMCs were acquired on a BD Fortessa X-20 on Day 0 and Day 5 of the stimulation. Cells were first stained with Live/Dead nIR (ThermoFisher) using the protocol provided by ThermoFisher, followed by Fc Block (BD Bioscience) in a 1:100 ratio for 20 minutes to reduced non-specific binding. Finally, the cells were stained for 30 minutes with the antibody master mix found in **Table S1**. Cells were then washed of any non-binding antibody and left in staining media till acquisition on a BD Fortessa X-20 with appropriate compensation.

### Flow cytometry analysis

Flow cytometry data were analysed using R (version 4.2.3) with the *ggplot2, MASS*, and *flowCore* packages. Compensation and transformation were applied to standardise fluorescence intensity values and facilitate downstream gating. Raw flow cytometry .fcs files were imported and processed using *flowCore*. Compensation was applied to correct for spectral overlap between fluorophores. Fluorescence intensities were then transformed using a logicle transformation to accommodate the wide dynamic range of fluorescence signals while preserving negative values. The transformation was applied using the following parameters:

where w controls the width of the linearization region, t is the data range (e.g., 18-bit data), m specifies the number of decades in the logarithmic portion, and a represents the additional negative range. Gating was performed using fluorescence-minus-one (FMO) controls, ensuring precise identification of cell populations. Sequential gating was applied to exclude debris and doublets before identifying target immune cell subsets based on marker expression. As each biological sample was processed in triplicate, the data from triplicate runs were averaged for robustness. Statistical analyses were performed on these averaged values. Proportional change (Δ𝐶) in cell composition proportion of cell type X between days 0 and 5 were calculated as:

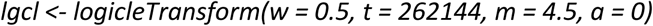

where *P^x^_day y_* is the proportion of cell type x on day y as a % of B cells.

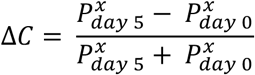

Proliferation modelling was carried out using the *nor1mix* package, which fits multi-Gaussian distributions to dye intensity histograms to infer cell division indices. The background population was first fitted using an unstimulated control. This baseline was then applied to model the division profile of stimulated samples. From the fitted models, proliferation index (PI), division index (DI), and percent divided were extracted for quantitative comparison between conditions. The division index (DI) was calculated as the average number of B cell divisions across the stimulated (Day 5) sample. The proliferation index (PI) calculated as the average number of B cell divisions excluding the undivided population.

All statistical analyses and visualisation were performed using base R plotting functions. Significance between conditions for BCR and FACS was assessed using MANOVA or non-parametric equivalents as appropriate.

### Single cell analysis

Peripheral blood single cell multi-omics data from COMBAT^72^ was extracted, including gene expression, cell-surface protein expression and VDJ BCR sequencing. Naïve B cells from the 10 healthy controls were extracted from this object, and only individuals with >100 naïve cells were retained (from 8 individuals. This resulted in a dataset of 1,192 high-confidence naïve B cells. The proliferation score and BCR signalling score was calculated using the *AddModuleScore* function in *Seurat* using the GOBP_B_CELL_PROLIFERATION and KEGG_B_CELL_RECEPTOR_SIGNALING_PATHWAY gene sets, respectively.

Correlations between expression and/or proliferation/BCR signalling scores were modelled as Mixed-Effects Model treating patient as a random effect using lme4 (given that multiple measurements per patient such as in scMulti-omics data). P-values were extracted using Satterthwaite’s approximation using lmerTest and standardized regression coefficient (β*) where beta_std ≈ partial correlation between X and Y adjusted for patient using fixef() function in lme4.

## Code and data availability

All code is available via https://github.com/rbr1/FMS_B_cell_tolerance. Data will be made available via EGA (currently in progress). Flow cytometry data will be made available upon request to authors.

## Acknowledgements

Firstly, we would like to thank the patients and clinicians who contributed to this study including our patient research partners. R.J.M.B.-R. and F.A.T. were supported by the University of Oxford. R.J.B is supported by a Versus Arthritis Fellowship (22976) and by a PhD grant from the Pain Relief Foundation. The authors acknowledge the Liverpool University Biobank, Faculty of Health and Life Sciences, University of Liverpool for some of the biological material.

## Authors’ contributions

A.G. and R.B-R conceived and designed the analysis. A.L., A.C.C., O.O., R.B., D.P., J.P., K.P., H.N., and F.T. collected the data. A.L., A.C.C., O.O., R.B., A.C, F.T., R.B-R contributed data or analysis tools. A.L., A.C.C., O.O., R.B., R.B-R performed the analysis. A.L., A.C.C., O.O., R.B., F.T., A.G., R.B-R contributed intellectual input/interpretation. A.L., A.C.C., O.O., R.B., F.T., A.G., R.B-R wrote the paper. All authors reviewed the paper.

## Declaration of interests

R.J.M.B.-R. is a co-founder of Alchemab Therapeutics Ltd and consultant for Alchemab Therapeutics Ltd, Roche, Enara Bio, UCB and GSK. A.G. has received consultancy fees from UCB. R.J.B has received consultancy fees from UCB and BioHaven. D.P. is an employee of Immunocore Ltd.

## Ethics approval and consent to participate

Informed consent was obtained for all patients. The study was in strict compliance with all institutional ethical regulations.

---

**Supplemental Figure 1.**
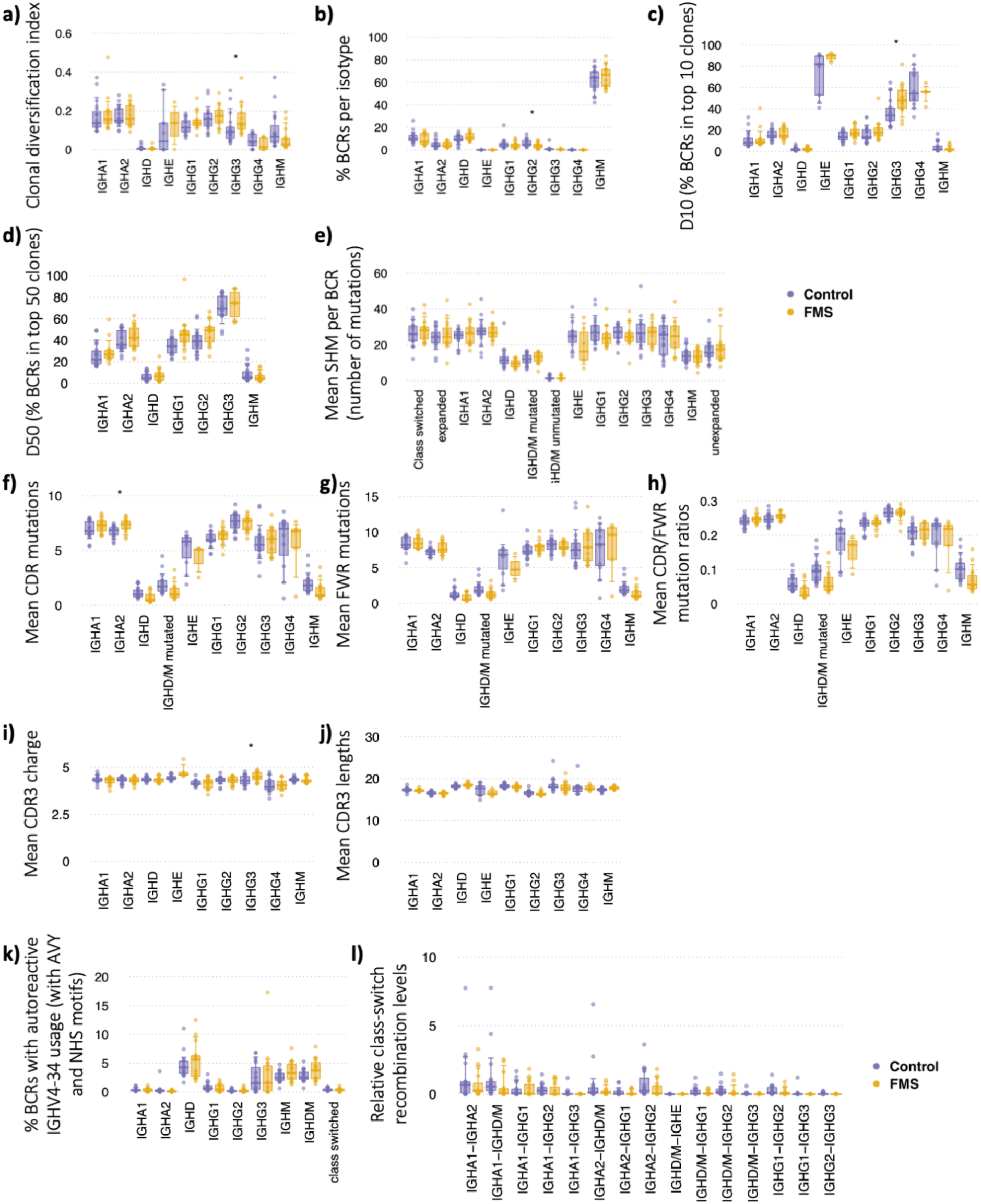
Boxplots of **a)** clonal diversification index, **b)** percentage BCRs per isotype, **c)** D10 (percentage BCRs comprising top 10 clones), **d)** D50 (percentage BCRs comprising top 50 clones), **a) e)** mean SHM per BCR, **f)** mean SHM within complementary determining regions (CDR) of the BCR, **g)** mean SHM within framework regions (FWR) of the BCR, **h)** mean ratio of FWR/CDR mutations, **i)** mean CDR3 charge, **j)** mean CDR3 lengths, **k)** percentage of IGHV4-24 BCRs containing the autoreactive NHS and AVY motifs, and **l)** relative CSR levels, stratified by isotype. Statistical testing used two-sided MANOVA; * denotes p < 0.05, ** denotes p < 0.005. Boxplots represent the 10th, 25th, 50th (median), 75th, and 90th percentiles. FMS: n = 14; Healthy Controls: n = 14.

**Supplemental Figure 2.**
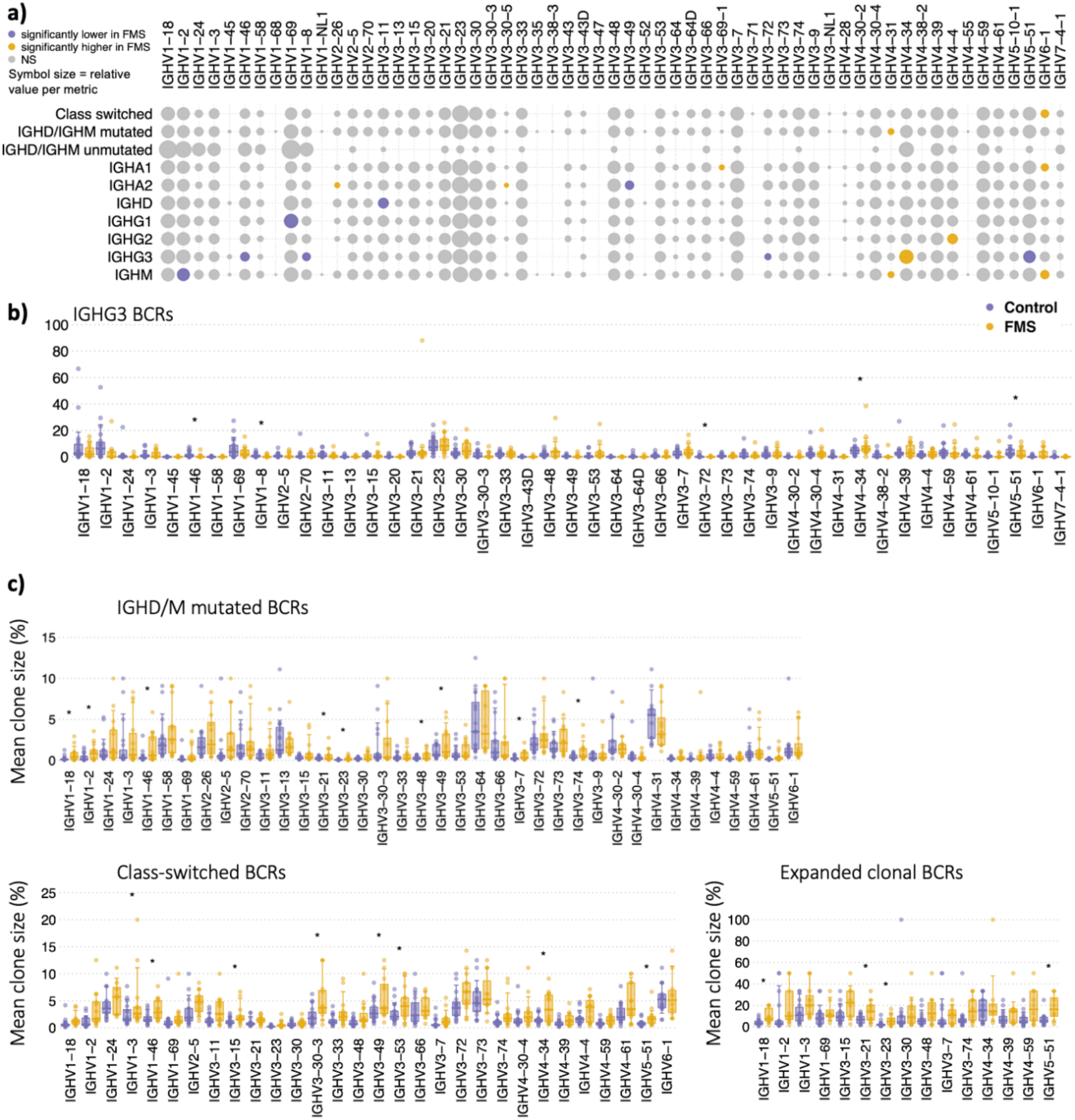
**a)** Heatmap of IGHV gene usage frequencies, stratified by isotype, and also by unmutated IGHM/D (representing predominantly naïve B cells), and mutated IGHM/D and class-switched BCRs (representing antigen experienced B cells). Yellow circles indicate significantly elevated frequencies in FMS; purple circles denote enrichment in controls. Circle size reflects the relative mean difference. **b)** Boxplots of percentage IGHG3 IGHV gene usages. **c)** Boxplots of mean clone sizes per IGHV gene split by isotype group. Only IGHV genes are plotted with >10 reads per patient for more than 10 patients. Statistical testing used two-sided MANOVA; * denotes p < 0.05, ** denotes p < 0.005. Boxplots represent the 10th, 25th, 50th (median), 75th, and 90th percentiles. FMS: n = 14; Healthy Controls: n = 14.

**Supplemental Figure 3.**
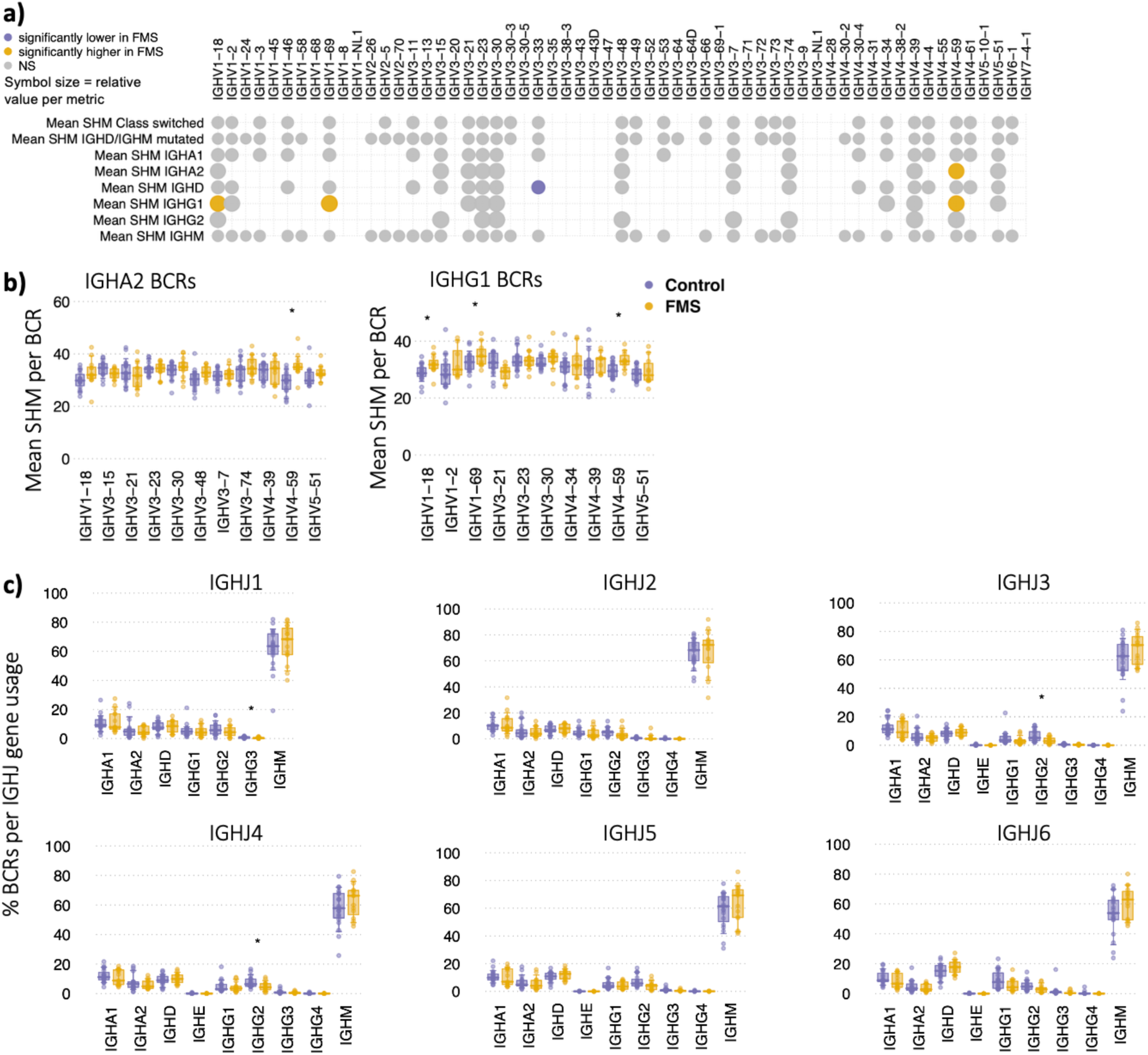
**a)** Heatmap of mean IGHV gene SHM levels, stratified by isotype, and also by unmutated IGHM/D (representing predominantly naïve B cells), and mutated IGHM/D and class-switched BCRs (representing antigen experienced B cells). Yellow circles indicate significantly elevated frequencies in FMS; purple circles denote enrichment in controls. Circle size reflects the relative mean difference. **b)** Boxplots of mean SHM per IGHV gene for IGHA2 and IGHG1. **c)** Boxplots of the IGHJ gene percentage usages stratified by isotype. Statistical testing used two-sided MANOVA; * denotes p < 0.05, ** denotes p < 0.005. Boxplots represent the 10th, 25th, 50th (median), 75th, and 90th percentiles. FMS: n = 14; Healthy Controls: n = 14.

**Supplemental Figure 4.**
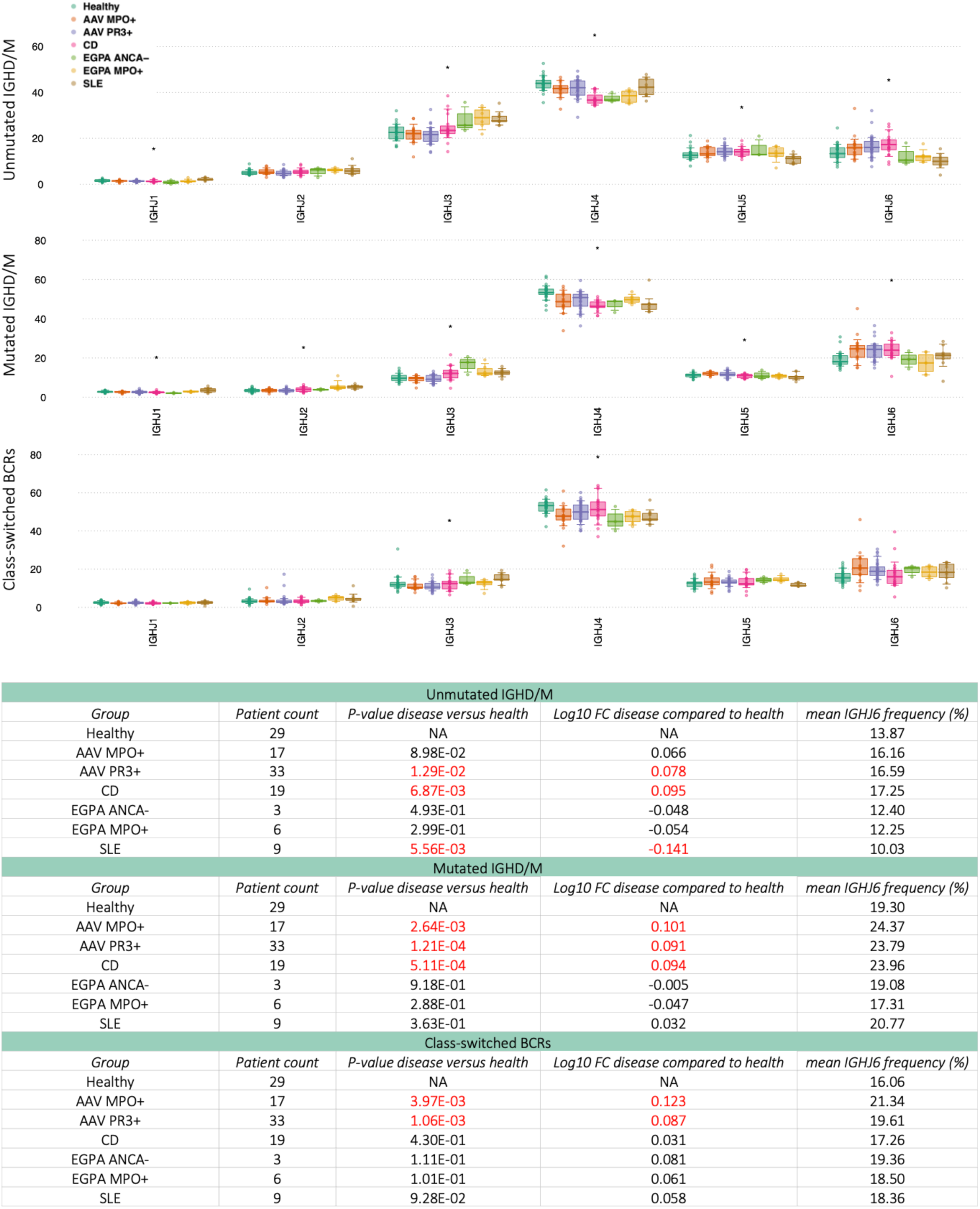
Reanalysis of the BCR repertoires from ^42^, (top) boxplots of the IGHJ gene usages stratified by IGHD/M and SHM status and (bottom) statistical analyses between healthy patients and disease groups. Statistically significantly different comparisons are highlighted in red.

**Supplemental Figure 5.**
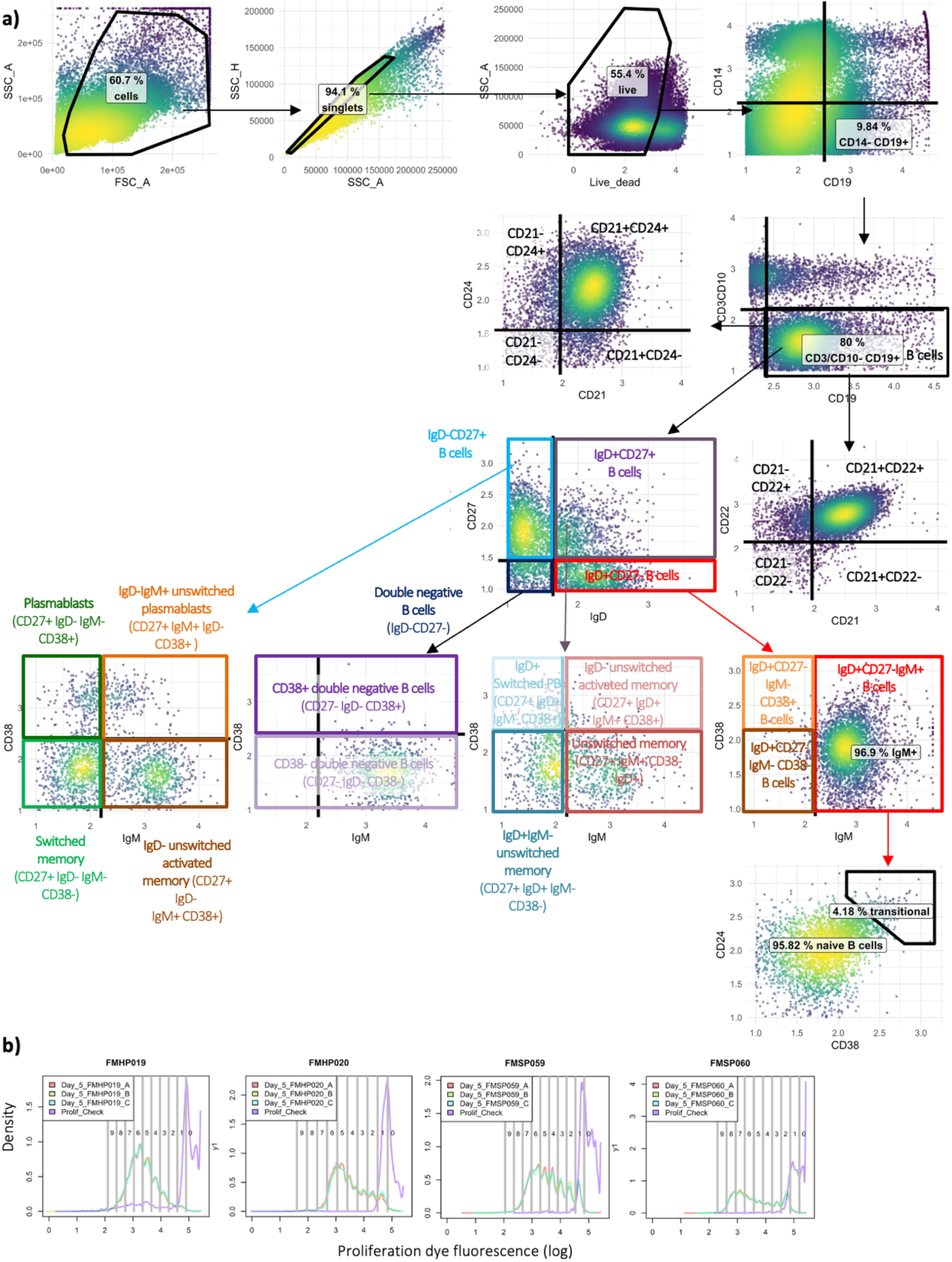
**a)** The flow cytometry gating strategy to define B cell subsets. **b)** Representative proliferation modelling plots for two healthy control patients (left) and two FMS patients (right).

**Supplemental Figure 6.**
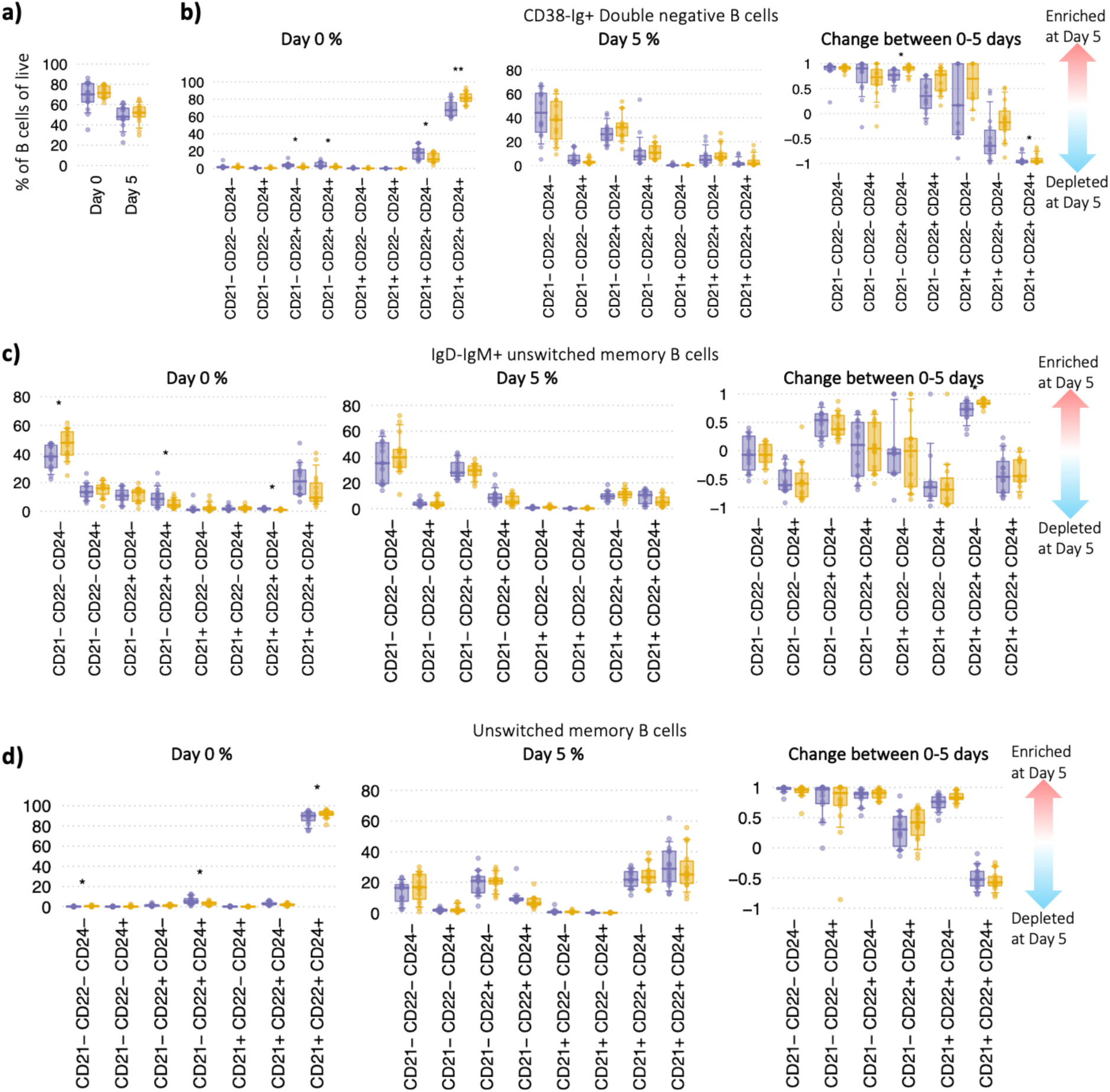
**a)** Proportion of total CD19⁺ B cells among live PBMCs at day 0 and day 5. **b-d)** Boxplots showing the day 0, day 5 and proportional changes in B cell subset composition by CD21/CD22/CD24 combination for **b)** CD38-IgM?+ Double negative B cells, **c)** IgD-IgM+ unswitched memory B cells, and **d)** Unswitched memory B cells, as a proportion of their subset. Statistical testing used two-sided MANOVA; * denotes p < 0.05, ** denotes p < 0.005. Boxplots represent the 10th, 25th, 50th (median), 75th, and 90th percentiles. FMS: n = 14; Healthy Controls: n = 14.

**Supplemental Figure 7.**
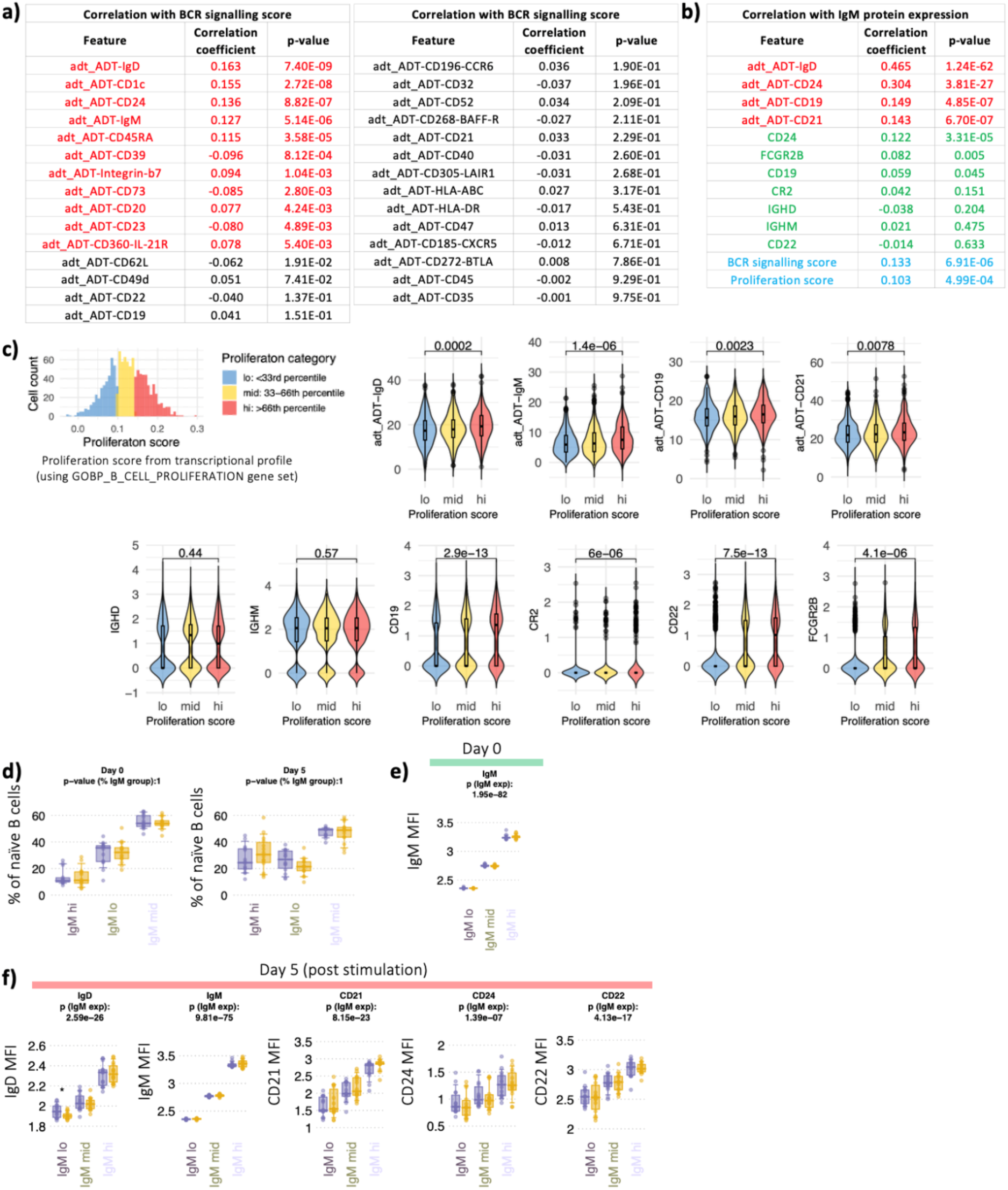
**a-c)** Re-analysis of single-cell multi-omic data from COMBAT using 8 healthy controls, high-confidence naïve B cells. **a)** Correlation of all highly expressed cell-surface proteins with BCR signalling score. **b)** Correlation of cell-surface protein expression (green), gene expression (red) and proliferation score (blue) with IgM protein expression. **c)** Naïve B cells were divided into the lowest, mid and highest proliferation score thirds and correlated these cell populations with surface IgD, IgM CD19 and CD21 protein expression, and IGHD, IGHM, CD19, CR2 (CD21), CD22 and FCGR2B gene expression via cell violin plots. **d)** Boxplots of the proportion of naïve B cell within the into IgM-lo, IgM-mid, and IgM-hi naïve B cell gates for healthy controls and FMS at day 0 and day 5. **e)** Distribution of IgM expression (mean fluorescence intensity, MFI) in naïve B cells, stratified into IgM-lo, IgM-mid, and IgM-hi naïve B cell subsets for day 0. **f)** Surface expression levels of IgD, IgM, CD21, CD22, CD24 of FMS and healthy controls within IgM-lo, -mid, -hi naïve B cells, grouped by patient group on day 5. P-values at the top of each plot gives the overall correlation of each marker with IgM expression. Statistical testing for (a)-(c) provided in methods Statistical testing for (d)-(f) used two-sided MANOVA; * denotes p < 0.05, ** denotes p < 0.005. Boxplots represent the 10th, 25th, 50th (median), 75th, and 90th percentiles. FMS: n = 14; Healthy Controls: n = 14.

**Supplemental Figure 8.**
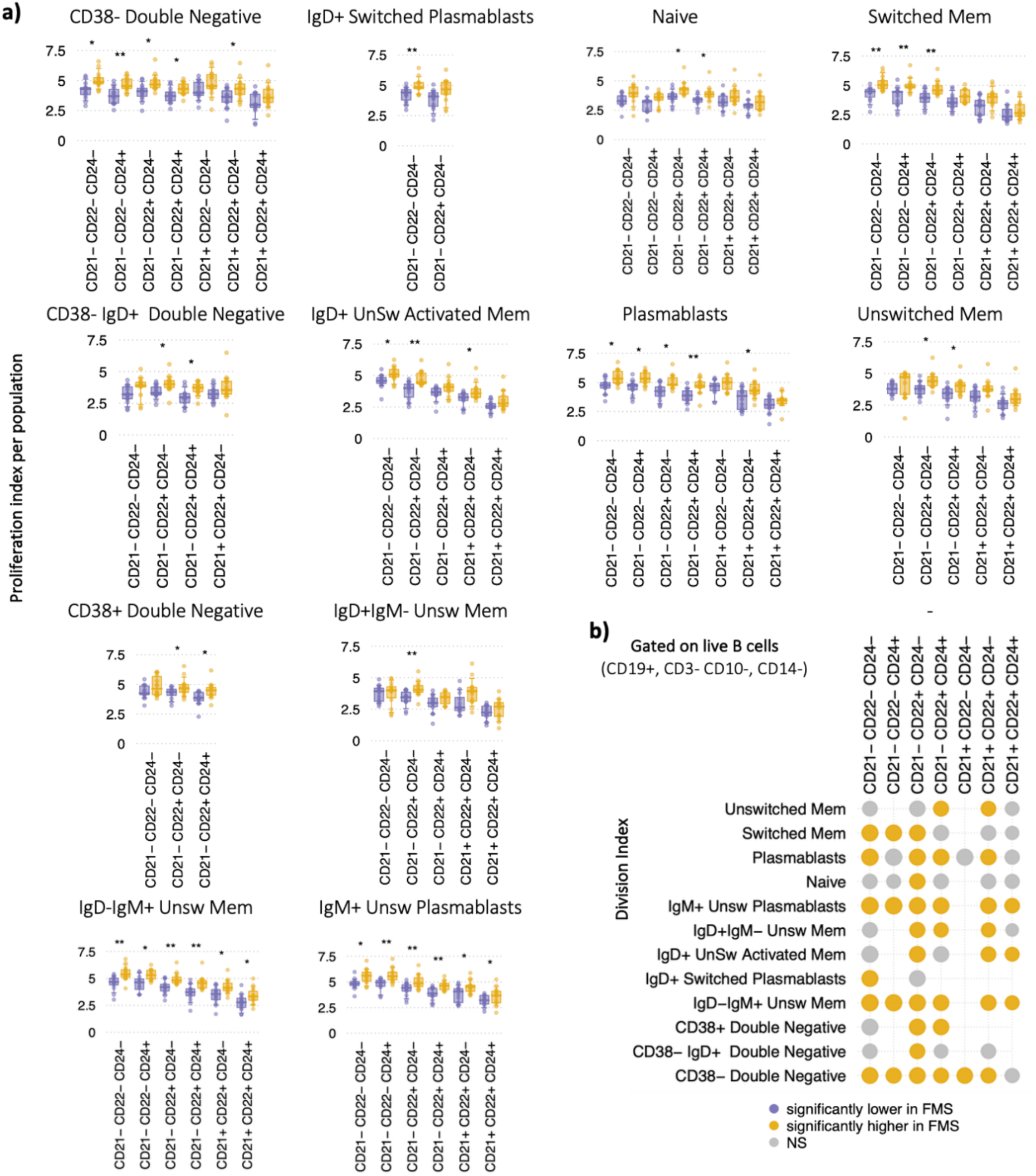
**a)** Boxplots of the proliferation indices per patient sample across all B cell subsets, stratified by CD21, CD22 and CD24 levels. IgM MFI between naïve IgM-lo, IgM-mid and IgM-hi B cells. **b)** Heatmap summarising differences in the division index after 5 days of *ex vivo* stimulation, stratified by CD21, CD22, and CD24 surface marker across B cell subsets. Yellow circles indicate significantly elevated frequencies in FMS; purple circles denote enrichment in controls. Circle size reflects the relative mean difference. Statistical testing used two-sided MANOVA; * denotes p < 0.05, ** denotes p < 0.005. Boxplots represent the 10th, 25th, 50th (median), 75th, and 90th percentiles. FMS: n = 14; Healthy Controls: n = 14.

